# The Coding sequence of firefly luciferase reporter gene affects specific hyperexpression *Arabidopsis thaliana cpl1* mutant

**DOI:** 10.1101/145011

**Authors:** Hisashi Koiwa, Akihito Fukudome

## Abstract

Forward genetic screening of mutants using firefly luciferase (*LUC*) reporter gene became a standard practice in plant research. Such screenings frequently identified alleles of *CPL1* (Carboxyl-terminal Phosphatase-Like 1) regardless of promoters or pathways studied. Expression of the corresponding endogenous genes often shows the minimal difference between wild type and *cpl1.* Here we show that the *LUC* coding sequence is responsible for the high expression in *cpl1,* using a classical *RD29a-LUC*. Deletion of the *LUC* 3’-UTR did not change hyperactivation of *LUC* in *cpl1.* However, a codon-modified *LUC* (*LUC2*) produced similar expression levels both in wild type and in *cpl1*. These results indicate that the coding region of *LUC* is responsible for the *cpl1*-specific *LUC* overexpression uncoupled with the expression of the endogenous counterpart.

---

Use of the reporter genes to monitor gene expression has become a standard practice in the characterization of genes of interest. Various reporter genes with different benefits include bacterial β-glucuronidase (GUS), fluorescent proteins from jellyfish/coral organisms (XFPs), and luciferase from bacteria (LUX), firefly (LUC) and sea pansy (*Renilla* LUC). Of these, LUC from firefly has been widely used to conduct non-invasive monitoring of plant gene expression. Due to the short half-life of *LUC* mRNA (45 min) and protein (15.5 min with luciferin, 155 min without), LUC provides a low-background and highly sensitive way to monitor the plant gene expression in real time and to study the inducible gene expression in response to environmental stimuli ^1^. Several groups including ours took advantage of LUC system to identify genetic mutations in *Arabidopsis thaliana,* where *LUC* reporter lines were subjected to mutagenesis, and genetic mutations were identified based on the alteration of LUC expression profile ^2^. Typically, inducible promoters were fused to LUC, and plants that over- or under-express LUC upon stimulation were identified as potential mutants for regulators of gene expression.

Over the past years, it became evident that the *LUC* reporter-based forward genetic approaches repeatedly identify mutations in *CPL1/FRY2* as well as *HOS5/RCF3* genes regardless of the biological processes studied ^3-8^. These include salt/osmotic stress, low-temperature-stress, jasmonate signaling/wounding, miRNA expression. Interestingly, in these studies, *cpl1/fry2* and *hos5/rcf3* mutations typically yield 10-100-fold enhancement in LUC mRNA or activities, however, the expression of corresponding endogenous transcripts was enhanced only ∼2 fold. Because endogenous stress-responsive transcripts are often induced >10 fold, the impact caused by the mutations are relatively subtle. The typical explanation for this inconsistency was the likely presence of additional negative regulatory elements in the endogenous promoter, which was not included in the promoter fragments used in the reporter constructs.

*CPL1/FRY2* encodes a protein with phosphatase activity that resembles eukaryotic TFIIF-interacting carboxyl terminal phosphatase (FCP1) and can dephosphorylate RNA polymerase II in vitro ^3, 4, 9^. As oppose to the expected function of CPL1 in the regulation of RNA polymerase II transcription, however, in vivo data are indicative that CPL1 functions in post-transcriptional metabolism of mRNA ^5, 7^. The first reported evidence that CPL1 affects the state of LUC reporter mRNA was that a *cpl1* mutant showed higher capping efficiency and altered polyadenylation site of *LUC* mRNA. Another study demonstrated that CPL1/FRY2 regulate RNA decay pathway of abnormal transcripts, such as transcripts that retain one or more introns ^10^. These findings prompted me to re-investigate whether stress-inducible promoter/signaling or reporter gene structure are the determinants of the enhanced expression of stress-inducible reporter gene in the *cpl1* mutant.

In addition to *RD29a-LUC* used in previous studies, we prepared *RD29a-LUCΔ3’* and *RD29a-LUC2*. *RD29a-LUCΔ3’* was made by removing 3’-UTR sequence of original firefly *LUC* mRNA from *RD29a-LUC*. The 3’-UTR of *LUC* mRNA contains two cryptic polyadenylation sites, usage of which were affected in *cpl1* mutants ^5^. On the other hand, *RD29a-LUC2* used codon-modified LUC (LUC2, Promega) coding sequence without firefly *LUC* 3’-UTR. These reporter cassettes were transformed into *Arabidopsis thaliana* Col-0 *cpl1-6* mutant line, and then wild-type lines were prepared by backcrossing T1 plants showing 3:1 segregation ratio to the wild-type Col-0 plants. Homozygous T3 (*cpl1-6*) and F3 (wild type) lines were used to directly compare the reporter gene expression levels between wild type and the *cpl1* mutant.

As shown in Figure 1, the *cpl1-6* mutant host strongly enhanced *RD29a-LUC* as reported previously. Removal of *LUC* 3’-UTR did not alter hyper-induction of *LUC* in *cpl1-6,* suggesting *LUC* hyperexpression in *cpl1-6* was not due to cryptic polyadenylation sites in *LUC* 3’-UTR. Notably, overall expression of *RD29a-LUCΔ3’* was enhanced compared to *RD29a-LUC.* Expression of codon-modified *RD29a-LUC2* substantially altered reporter gene expression profile. Compared to *RD29a-LUC, RD29a-LUC2* activity was similarly increased in both wild type and *cpl1-6*. The induction pattern was close to that of endogenous *RD29a* mRNA level measured by RT-qPCR (Figure 1C), albeit *RD29a-LUC2* response was slower than that of *RD29a* mRNA. This result suggested that hyperexpression of *RD29a-LUC* in *cpl1* mutant are neither due to the enhancement of *RD29a* promoter activity, upstream osmotic-stress signaling events nor alternative polyadenylation in 3’-UTR. Instead, the data showed that *LUC* coding sequence strongly influences *cpl1*-dependent hyperexpression. This reporter-gene-specific effect by the cpl1 mutation explains why mutations in *CPL1/FRY2* as well as in *HOS5/RCF3* encoding a binding partner of CPL1 are repeatedly identified in *LUC* reporter gene-based forward genetic screening, and why there are discrepancies between *LUC* expression level and endogenous counterpart in *cpl1/fry2*. Because LUC mRNA has a short half-life in the plant cell ^1, 11^, the likely mechanism is *LUC* mRNA but not endogenous counterpart being a target of CPL1-dependent RNA decay pathway ^10^. This observation suggests that we need to be cautious about the impact of *cpl1* on synthetic phenotype produced by artificial transgenes, because *cpl1* may produce phenotype by specifically affecting the level of transgene expression.

**Figure 1.**
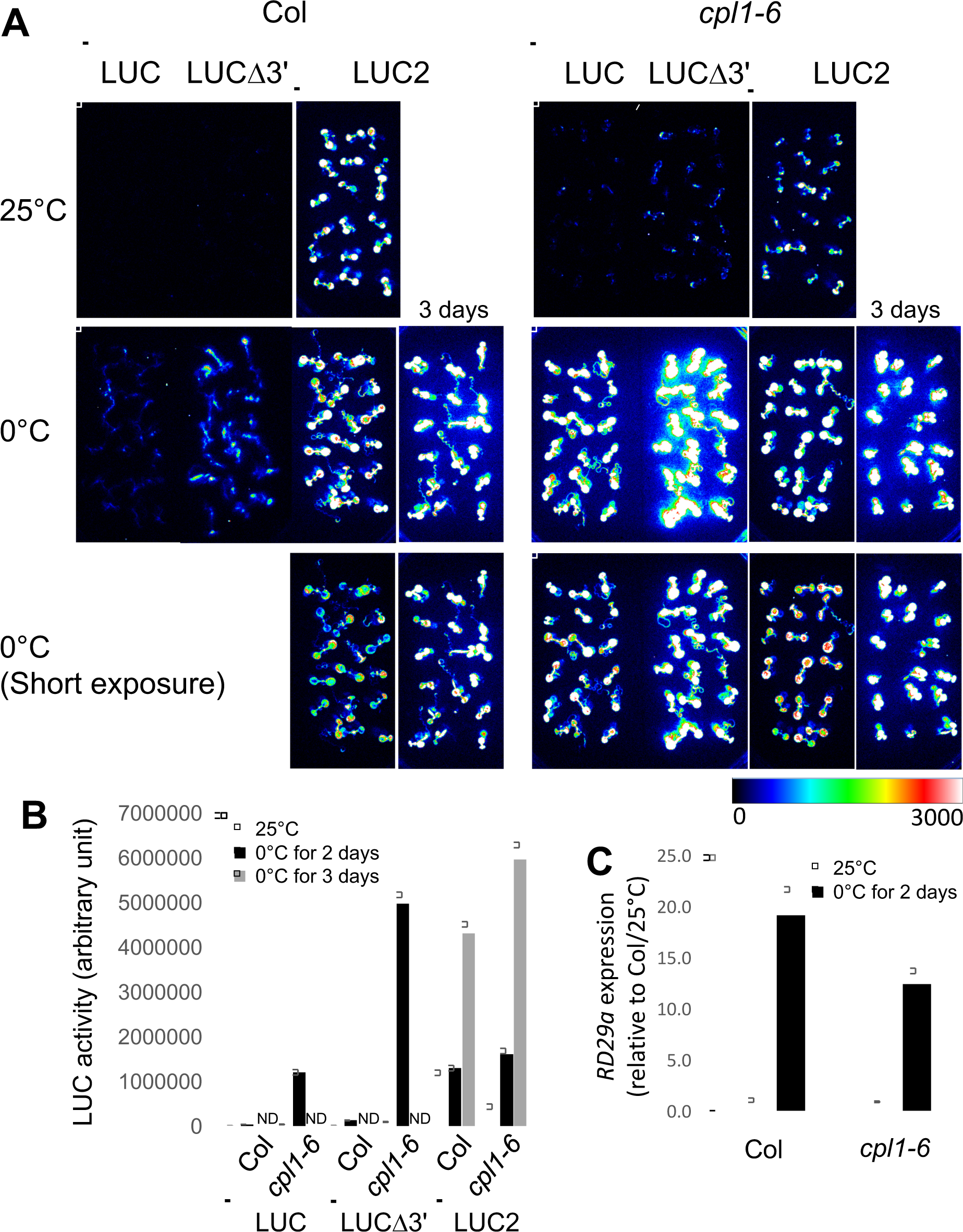
Expression of *RD29a*-luciferase reporter genes in Col-0 and *cpl1-6.* **A)** Bioluminescence images of *RD29a-LUC* WT, Δ3’, and *LUC2* in 25°C and after 2 or 3-day 0°C treatment. Plants were grown on medium containing 1/4x MS salts, 1% sucrose, and 0.7% agar for 7 days. For cold treatment, plates were incubated at 0°C in the dark for indicated duration. Exposure time for CCD camera was 10 min (5 min for short exposure panels). **B**) Bioluminescence image quantification results. Each data point was calculated using 50-70 plants from 3-4 independent plates. **C**) RT-qPCR analysis results of *RD29a* transcripts in Col-0 and *cpl1-6.* Total RNA was extracted from plants prepared the same way as A and B.

I found the expression of codon-modified *LUC2* reporter gene retains stress-regulation in wild type and is much less affected by *the cpl1-6* mutation, therefore, *LUC2* provides a promising alternative for reporter-based forward-genetic screening. It should be noted, however, that *RD29a-LUC2* produced higher background than RD29a-LUC, and there was a delay in the timing of induction of *RD29a-LUC2* activity above the background level after the onset of cold treatment relative to *RD29a-LUC*, which would require careful optimization of the assay system.

## Materials and methods

### Preparation and analysis of reporter lines

*RD29a-LUCΔ3’* was prepared by inserting a PCR fragment containing 3’-end of *LUC* coding sequence (primer pair 1358 TAGTTGACCGCTTGAAGTCT/1359 GGAAAAGAGCTCTTACAATTTGGACTTTCCGC) into PacI-SacI sites of original *RD29a-LUC* plasmid ^2^. *RD29a-LUC2* was prepared by inserting a PCR fragment encoding *LUC2* (primer pair: 1471 CTATTTACAATTACAGTCGACATGGAAGATGCCAAAAACATT/1472 cagatcccccgggggtaccgagctcTTACACGGCGATCTTGCCGC) into SalI-SacI site of *the RD29a-LUC* plasmid. Resulting plasmids were introduced into *Agrobacterium tumefaciens* ABI, and used for transformation of *cpl1-6* ^12^ by flower transformation. Kanamycin-resistant plants were selected as described ^13^. Transformants whose luciferase expression phenotype were segregating with 3:1 ratio were backcrossed to Col-0. Homozygous T3 (*cpl1-6* background) and F3 (Col-0 background) lines were established for each transgene and were used for the expression analysis. Luciferase imaging analysis and RT-qPCR analysis was performed as described ^12^.

## Acknowledgement

This study was partially supported by USDA-NIFA (2015-67013-22816).

## Notes

Conflict of interest: There is no conflict of interest for this study.

